# DNA unwinding is the primary determinant of CRISPR-Cas9 activity

**DOI:** 10.1101/205823

**Authors:** Shanzhong Gong, Helen Hong Yu, Kenneth A. Johnson, David W. Taylor

## Abstract

Bacterial adaptive immunity utilizes RNA-guided surveillance complexes composed of CRISPR (clustered regularly interspaced short palindromic repeats)-associated (Cas) proteins together with CRISPR RNAs (crRNAs) to target foreign nucleic acids for destruction. Cas9, a type II CRISPR-Cas effector complex, can be programed with a single guide RNA that base-pairs with the target strand of dsDNA, displacing the non-target strand to create an R-loop, where the HNH and RuvC nuclease domains can cleave opposing strands. Cas9 has been repurposed for a variety of important genome engineering applications. While many structural and biochemical studies have shed light on the mechanism of Cas9 cleavage, a clear unifying model has yet to emerge. Our detailed kinetic characterization of the enzyme reveals that DNA binding is reversible, R-loop formation is rate-limiting, occurring in two steps, one for each of the nuclease domains. Although the HNH nuclease activity is stimulated by Mg^2+^ with a single measureable *K*_*d*_, the RuvC activity requires two distinct Mg^2+^ binding events. The specificity constant for cleavage is determined through an induced-fit mechanism as the product of the equilibrium binding affinity for DNA and the rate of R-loop formation.

## Introduction

Bacteria and archaea defend themselves against infection using RNA-guided adaptive immune systems comprising CRISPR (clustered regularly interspaced short palindromic repeats) loci and their associated *cas* genes (Wiedenheft et al., 2012, Sorek et al., 2013, Barrangou et al., 2007, Marraffini and Sontheimer, 2008). Small fragments (or spacers) of foreign DNA are integrated into the CRISPR repeat-spacer array within the host chromosome as new spacers during acquisition (Amitai and Sorek, 2016), providing a memory of infection. Transcription and processing of the CRISPR array produce mature CRISPR RNAs (crRNAs) that associate with the Cas protein effector complexes (Brouns et al., 2008). A hallmark of CRISPR–Cas systems is the assembly Cas proteins around crRNAs to produce RNA-guided surveillance complexes to interrogate DNA targets and destroy matching sequences in foreign nucleic acids (Wiedenheft et al., 2012, van der Oost et al., 2014). Base-pairing between the crRNA and a complementary foreign target sequence (protospacer) triggers sequence-specific degradation of targets after the protospacer adjacent motif (PAM) recognition (Garneau et al., 2010, Gasiunas et al., 2012, Marraffini and Sontheimer, 2008). The PAM, a ~3-5-bp motif located in close proximity to the target sequence, is important in target identification and licenses the crRNA-effector complexes for cleavage after recognition (Mojica et al., 2009).

Six distinct types (I-VI) of CRISPR-Cas systems have been identified and each employs a unique set of Cas proteins for CRISPR-mediated interference (Makarova et al., 2015). While type I and type III crRNA-guided surveillance complexes are composed of multiple proteins that form larger complexes (van der Oost et al., 2014), type II complexes use a single endonuclease, Cas9, to recognize and cleave dsDNA substrates (Jinek et al., 2012). Both type I and type II CRISPR-Cas systems require R-loop formation, in which the guide RNA segment of crRNAs invades double-helical DNA to form an RNA-DNA hybrid helix with the target DNA strand while displacing the opposing non-target strand, for DNA degradation (Jore et al., 2011, Szczelkun et al., 2014, Rutkauskas et al., 2015, Blosser et al., 2015). In type II CRISPR-Cas systems, the Cas9 endonuclease creates an R-loop, where the target and non-target strands are funneled into and cleaved by the HNH and RuvC domains, respectively. R-loop generation and target cleavage by Cas9 requires both a crRNA and a trans-activating crRNA (tracrRNA) (Jinek et al., 2012). The crRNA and tracrRNA can be fused to from a chimeric single-guide RNA (sgRNA), which retains the ability to guide Cas9-mediated target degradation (Jinek et al., 2012).

Cas9 programmed with sgRNAs can be used to target specific DNA sequences in the genome to create a blunt-ended double-strand break (DSB) (Jinek et al., 2012). These DSBs are repaired by the host machinery to create random insertions and/or deletions (indels) or precise genome modification using a homologous template at the site of the DSB via non-homologous end-joining or homology-directed repair, respectively (Lieber, 2010, San Filippo et al., 2008). Cas9 has rapidly become a revolutionary tool for genome engineering in a wide range of organisms and systems because of its programmability, high efficiency, and ease of use (Hsu et al., 2014).

The general mechanisms behind Cas9 target recognition, R-loop formation, and cleavage activities have rapidly unfolded over the past several years. Crystal structures of Cas9 alone show it has a bi-lobed architecture, containing an alpha-helical lobe and a nuclease lobe (Jinek et al., 2014). Electron microscopy reconstructions of Cas9 bound to guide RNA or to DNA revealed substantial conformational rearrangements of Cas9 to form a DNA recognition-competent structure (Jinek et al., 2014). Later, crystal structures bound to a single-stranded target DNA provided further details into these structural changes, including PAM recognition (Nishimasu et al., 2014, Anders et al., 2014, Nishimasu et al., 2015). Notably, the crystal structure of Cas9 bound to guide RNA showed that the seed sequence of the RNA is pre-ordered to accommodate the target strand, and the majority of the reorganizations seen in the DNA-bound state have already occurred (Jiang et al., 2015). In all of these structures, the HNH domain is in a position that is incompatible for target strand cleavage. R-loop formation drives the repositioning and reorientation of the HNH domain into a cleavage-competent state as shown by a crystal structure of Cas9 with a partial R-loop and a cryo-EM reconstruction of a bona-fide R-loop structure (Jiang et al., 2016). Additionally, recent studies have suggested a strong correlation between R-loop stability and CRISPR-Cas9 cleavage efficiency (Xu et al., 2017).

Importantly, all of these structural studies suggest that the HNH domain is in a dynamic equilibrium between open and closed conformations, with the closed conformation being required for target cleavage. Recently, molecular mechanisms governing the HNH and RuvC nuclease domains have begun to emerge. Bulk Forster Resonance Energy Transfer (FRET) assays have shown that there is a conformational activation that occurs upon HNH substrate docking that is important for RuvC-mediated cleavage. An alpha-helical linker region between the two nuclease domains acts as an allosteric switch between the repositioning of the HNH domain and the RuvC domain, activating RuvC for non-target strand cleavage (Sternberg et al., 2015). Single-molecule FRET experiments identified an intermediate state that occurs during HNH dynamics that prevents activation with mismatches (Dagdas et al., 2017). The RuvC domain of Cas9 shares structural similarity with *Escherichia coli* RuvC and *Thermus thermophilus* RuvC (Nishimasu et al., 2014), which cleaves the phosphodiester backbone through a two-metal ion mechanism (Ariyoshi et al., 1994, Chen et al., 2013). In addition, the HNH domain of Cas9 shares structural similarity with phage T4 endonuclease VII (Endo VII), which processes substrate through a one-metal ion mechanism (Biertümpfel et al., 2007).

Although many structural details have emerged from these studies, a comprehensive or unifying model for the Cas9 cleavage pathway is lacking, in part, because many biochemical studies have relied on limited observations of the cleavage activity or indirect measurements. Kinetic analysis is necessary to establish the enzymatic pathway, which cannot be defined from measurements at equilibrium. To define a complete framework for Cas9, we performed a detailed kinetic analysis using both stopped flow and quench flow techniques on Cas9 cleavage of a perfectly matched target DNA. We show that R-loop formation and cleavage are rapid reactions, reaching completion in approximately 5 seconds. The rate-limiting step is R-loop formation, which appears to occur in two steps, one leading to HNH cleavage and a second preceding RuvC cleavage. We also show that HNH and RuvC nuclease-mediated cleavage can occur simultaneously, and appear to use one-and two-metal ion mechanisms, respectively. Finally, we document a nonproductive state accounting for a slow cleavage reaction for ~15% of the total enzyme.

## Results

### Cas9 is a single turnover enzyme that binds DNA cleavage products tightly

While many biochemical experiments have been performed on Cas9, none have reported an active site concentration, which is required to accurately assess the mechanism. To measure the active site concentration of *Streptococcus pyogenes* Cas9 (Cas9) programmed with sgRNA (Cas9.gRNA), a fixed concentration of enzyme (100 nM, estimated from absorbance at 280nm) of Cas9.gRNA was allowed to react with various concentrations of matched target DNA (5′-end labeled on the target strand) in the presence of 10 mM Mg^2+^. After 30 min at 37 °C, the products were resolved on a sequencing gel, quantified, and plotted as a function of DNA concentration (Figure 1A). These direct measurements established an active enzyme concentration of 28 nM. It is important to note that the concentration of DNA required to saturate activity was equal to the concentration of product formed, so the active site concentration is defined by the intersection of the two straight lines in Figure 1A. If a large fraction of inactive enzyme bound DNA but did not react, then a higher DNA concentration would have been required to saturate the active sites, relative to the concentration of product formed. These results suggest that a large fraction of the enzyme is completely inactive, if one believes the estimate of protein concentration based on the absorbance measurement, which may include systematic errors. We use the active site concentration (Figure 1A) for all subsequent studies.

**Figure 1.**
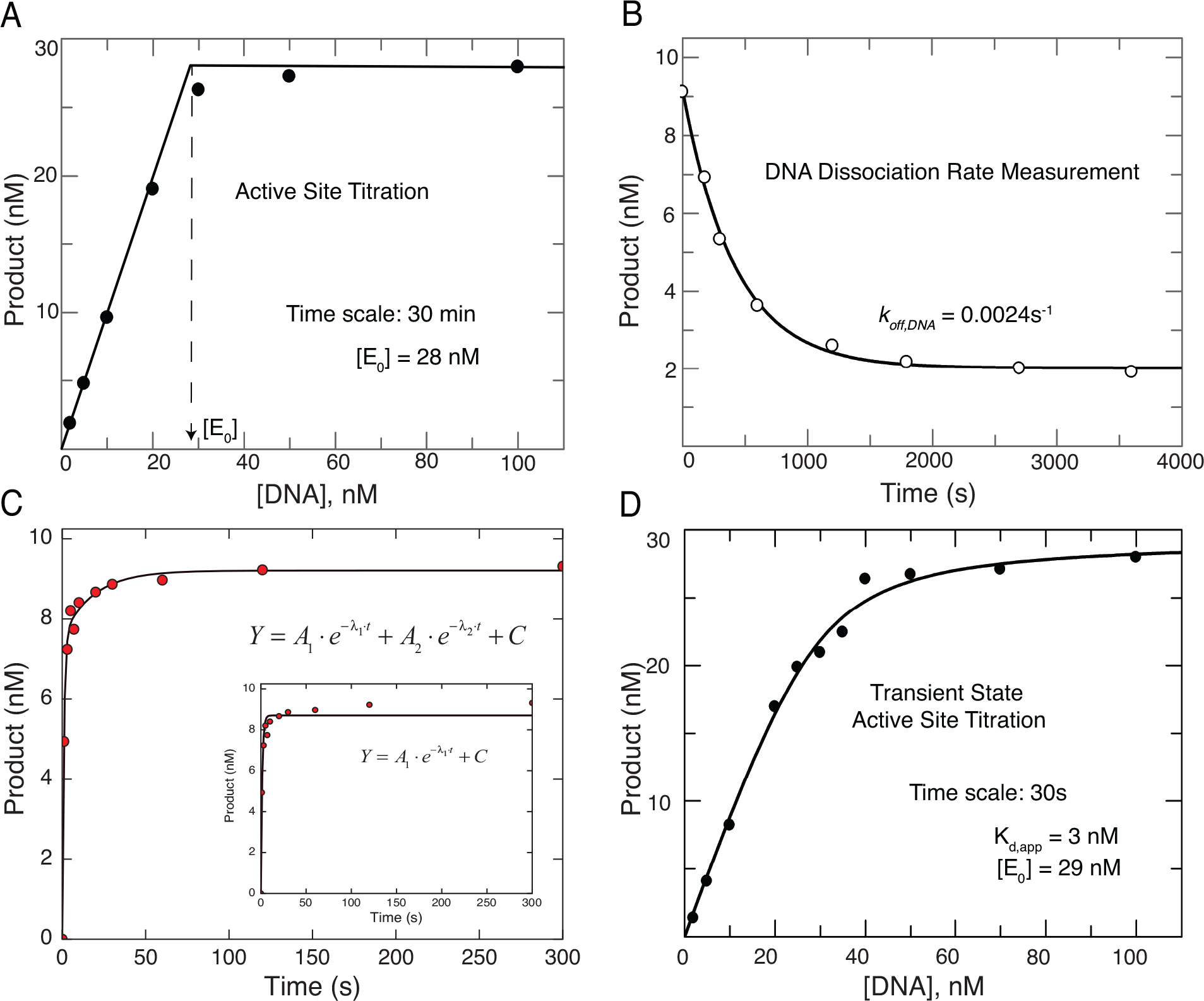
Cas9 is a single turnover enzyme that binds cleaved DNA products tightly. (A) Active site titration experiment was performed by mixing a fixed concentration (100 nM) of Cas9.gRNA (1:1 ratio) with variable concentrations of a perfectly matched γ-^32^P labeled DNA target. By allowing the reaction to reach completion (30 min), the results reveal the active site concentration of Cas9.gRNA. Two dotted lines shown in Figure 1A indicate the nearly irreversible binding curve; and vertical dotted arrow shown in Figure 1A indicates the active site concentration of Cas9.gRNA. (B) The DNA dissociation rate was measured by premixing γ-^32^P labeled perfectly matched target DNA (10 nM) and Cas9.gRNA (28 nM active site concentration) for 10 min in the absence of Mg^2+^ followed by addition of a large excess of unlabeled DNA trap (200 nM) for various incubation times (0, 1, 5, 10, 20, 30, 45, 60, 90 and 120 min), after which the chemical reaction was initiated by addition of 10 mM free Mg^2+^, and then quenched by addition of EDTA after 30 s. The amount of product formed versus time was fit to a single exponential decay equation to obtain the dissociation rate of DNA (*k*_*off,DNA*_ = 0.0024s^−1^). (C) Time course Cas9 cleavage assay performed by directly mixing γ-^32^P labeled DNA (10 nM) and Cas9.gRNA (28 nM active site concentration) in the presence of 10 mM Mg^2+^. Samples were then collected at different time points by stopping the reaction by adding EDTA. The concentration of product formed was fit to double exponential to account for the lower amplitude slow phase. A single exponential fit is shown in the inset. (D) The experiment in (A) was repeated but allowing after 30 s of reaction. The DNA concentration dependence of the amount of product formed provided an active site titration to define the apparent *K*_*d*_ for DNA binding (3 nM) as well as the active site enzyme concentration.

It is important to recognize that this experiment succeeds only because Cas9 is a single turnover enzyme as described previously (Sternberg et al., 2014). After each enzyme catalyzes a single round of cleavage reactions, it stops. Cas9 represents an extreme case of a pre-steady-state burst where product release is exceedingly slow. Moreover, because of the long time of incubation (as described below) and the irreversible cleavage reaction, the reaction proceeded to completion, and this precludes estimation of the *K*_*d*_ for DNA binding from the data in Figure 1A. As the reaction proceeds, the largely irreversible nuclease reaction pulls to completion any reversibly bound DNA binding until all of the DNA (excess enzyme) or active enzyme (excess DNA in this case) is consumed.

To investigate the extent to which DNA is reversibly bound before cleavage, we first measured the rate of DNA dissociation from a Cas9.gRNA.DNA complex in the absence of Mg^2+^. Cas9.gRNA was first incubated for 10 minutes with radiolabeled DNA in the absence of Mg^2+^ to allow formation of the Cas9.gRNA.DNA complex. We then added a 20-fold excess of unlabeled DNA as a trap and then after various times, the cleavage reaction was initiated by the addition of 10 mM Mg^2+^, allowed to proceed for 30 seconds, then quenched. The plot of product formed versus the time of incubation with the DNA trap was fit to single exponential decay equation to define the dissociation rate of DNA from Cas9.gRNA.DNA (*k*_*off,DNA*_ = 0.0024 s^−1^), giving a half-life of 5 minutes (Figure 1B). These results show that DNA dissociates slowly *prior to* target cleavage. This is not to be confused with a previous study that showed Cas9 does not dissociate from DNA *after* cleavage (Sternberg et al., 2014).

### Cas9.gRNA binds DNA reversibly prior to cleavage

To understand the reactions governing Cas9-mediated target cleavage, we need to monitor the reaction on the time scale of a single turnover of the catalytic reaction. We examined the time course by mixing Cas9.gRNA (28 nM active site) with 10 nM DNA in the presence of Mg^2+^. After quenching by adding EDTA, the products were resolved and quantified (Figure 1C). The time-dependence of HNH product formation curve was biphasic. In the initial fast phase, ~85% of the cleavage occurred at a rate of ~1 s^−1^, while the remaining 15% of the reaction occurred at a rate of ~0.03 s^−1^. The significance of the slow reaction will be described below.

Because the observed rate of cleavage (~1 s^−1^) is much faster than DNA dissociation (0.0024 s^−1^), we can now perform a transient state kinetic active site titration to measure the DNA binding affinity prior to cleavage. Because the reaction is faster than the time required for DNA binding to equilibrate, the amplitude of the rapid transient of DNA cleavage provides a direct measure of the concentration of the Cas9.gRNA.DNA complex formed and poised for catalysis after mixing. Cas9.gRNA was allowed to react with various concentrations of DNA (10 mM Mg^2+^). The concentration of product formed after 30s reaction was then plotted as a function of DNA concentration. Fitting this data to a quadratic equation gave a apparent *K*_*d*_ for DNA binding of 3 nM (Figure 1D). If the enzyme and DNA had been allowed to equilibrate before the reaction, this would represent the *K*_*d*_ for DNA binding. However, because of the very slow dissociation rate, this experiment provides an estimate of the relative rates of DNA binding versus cleavage to define the apparent *K*_*d*_.

### A minor nonproductive DNA binding state

The slow phase of the reaction (Figure 1C) suggests that ~15% of the DNA binds in a nonproductive state, or that 15% of the enzyme reacts more slowly. One possible explanation is that the slow phase represents a re-equilibration of the DNA binding, but this appears to be inconsistent with the DNA dissociation rate measured (Figure 1B). We first asked whether the slow phase was due to a nonproductive Cas9.gRNA.DNA complex, defined as a state where the DNA was tightly bound and reacted slowly without DNA dissociation. To test this postulate, a DNA trap was added simultaneously with Mg^2+^ to Cas9.gRNA.DNA, and the effect of the DNA trap on the slow phase of the cleavage reaction was observed (Figure 2A). There was a slight reduction in the amplitude of the slow phase in the presence of the DNA trap (red) compared to the absence (green). Thus at least part of the slow phase is due to weakly bound DNA such that dissociation competed kinetically with the rate of the chemical reaction.

**Figure 2.**
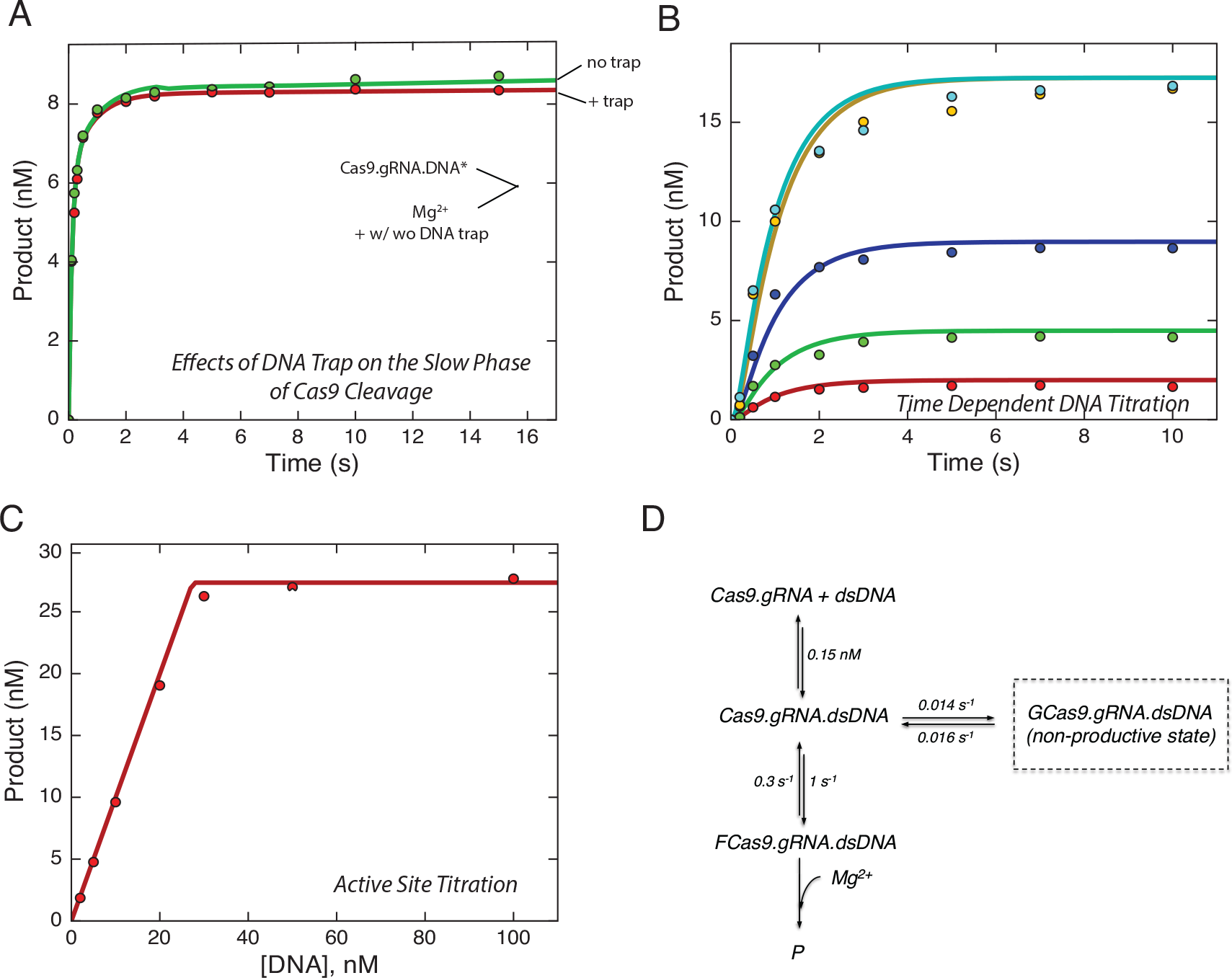
Cas9.gRNA exists in both a productive and nonproductive state for cleavage. (A) Effects of DNA trap on the slow phase of Cas9 cleavage was tested by premixing γ-^32^P labeled DNA (10 nM) and Cas9.gRNA (28 nM active site concentration) in the absence of Mg^2+^. The experiment was initiated by addition of 10 mM free Mg^2+^ in the presence and absence of DNA trap (200 nM unlabeled DNA substrate). (B) A time-dependence or reaction at various concentrations of DNA; the experiment was performed by mixing fixed a concentration of Cas9.gRNA (17.3 nM active site concentration) with variable concentrations of γ-^32^P labeled DNA (2, 5, 10, 20, 40 nM). Then, the reaction was quenched at varying time points by the addition of 0.5 M EDTA and the amount of product (HNH cleavage) was quantified. (C) Active site titration experiment as shown in Figure 1A. (D) The smooth lines in each of the above figures were derived from globally fitting to this model.

This analysis led to the suggestion that DNA binding may occur in several steps, with a weak initial binding followed by tighter binding. In order to test whether the slow phase seen in Figures 1C and 2B was due to a slow isomerization preceding the tight binding mode, we examined the kinetics of the reaction as a function of time by initiating the reaction by mixing Cas9.RNA with DNA in the presence of Mg^2+^. If the isomerization from weak to tight binding was a step on the pathway leading to catalysis, we should only observe the slow reaction phase. The time dependence of the reaction was tested at various concentrations of DNA (Figure 2B). In each case the reaction occurred at a rate of ~1 s^−1^ so it was completed in ~5 s. These data suggest that the slow phase of the reaction is due to a side reaction where the DNA binds in a non-productive state, either forming after DNA binding or due to an altered state of the enzyme existing before DNA binding. Global fitting of the above two experiments with the active site titration (Figure 2C) is indeed consistent with the formation of non-productive state of the enzyme. The transition from the nonproductive to the productive state of Cas9.RNA.DNA is much slower (0.016s^−1^) than catalysis by the majority of the enzyme (Figure 2D). For the remainder of this discussion, we will focus our attention on the faster reacting 85% of the enzyme.

### R-Loop formation is the rate-limiting step

We next sought to more accurately measure the kinetics of HNH and RuvC cleavage on a shorter time scale. In addition, we reasoned that we might see a difference depending on the starting conditions. If there is a slow step after DNA binding that limited the rate of cleavage, we might see a faster reaction if we preincubated the Cas9.gRNA with DNA in the absence of Mg^2+^ and then initiated the reaction by adding Mg^2+^ in contrast to simultaneously adding DNA and Mg^2+^ to the Cas9.gRNA. We performed parallel experiments using a DNA substrate 5′-end labeled on either the target-strand or non-target strand to measure cleavage rates for the HNH and RuvC nuclease domains, respectively.

First, we mixed Cas9.gRNA simultaneously with DNA and 10 mM Mg^2+^ to initiate the cleavage reaction. The plot of product formation as a function of time could be fit to a single exponential equation to define cleavage rates of 1s^−1^ and 0.2s^−1^ for the HNH and RuvC domains, respectively (Figure 3A). In the second experiment, we premixed Cas9.gRNA with DNA to form the Cas9.gRNA.DNA complex in the absence of Mg^2+^, and then the reaction was then initiated by the addition of 10 mM Mg^2+^. Again, a plot of cleavage product as function of time could also be fit to a single exponential equation to define cleavage rates, but the rates were faster: 4.3 s^−1^ and 3.5 s^−1^ for HNH and RuvC domains, respectively (Figure 3B). The observed cleavage rates of reactions initiated by addition of Mg^2+^ after Cas9.gRNA.DNA was allowed to equilibrate were significantly faster than the observed cleavage rates initiated by simultaneously mixing DNA and Mg^2+^ to Cas9.gRNA. Therefore, the rate-limiting step in the enzyme pathway is not the cleavage chemistry itself, but rather some isomerization of the Cas9.gRNA.DNA complex after DNA binding and preceding cleavage (Figure 3C).

**Figure 3.**
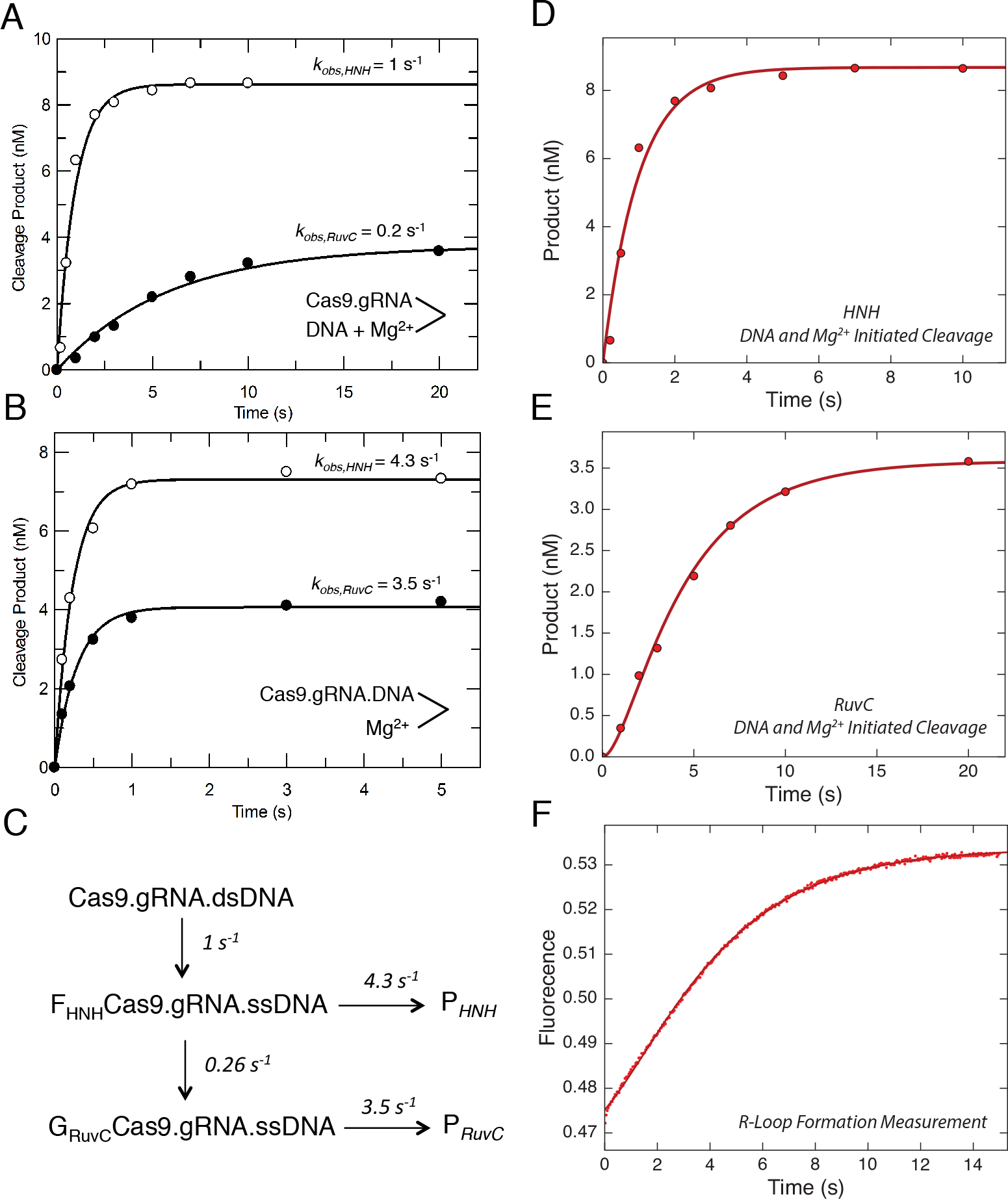
R-Loop formation is the rate-limiting step. (A) Cas9 cleavage rates were measured after the reaction was initiated by the simultaneous addition of DNA (10 nM) and Mg^2+^ (10 nM) to Cas9.gRNA (28 nM active site concentration). The reaction was quenched by the addition of EDTA at different time points, and then the results were fit to a single exponential equation. DNA was either labeled on the target strand or the non-target strand to obtain the cleavage rates from HNH (open symbols) and RuvC (closed symbols) nuclease activities, respectively. (B) HNH and RuvC cleavage rates measured after initiating the reaction by the addition of Mg^2+^ (10 nm) to a mixture of Cas9.gRNA (28 nM) with DNA (10 nm) pre-incubated for 10 min. Aside from the order of mixing, the experiment was performed and analyzed as in A. (C) Model showing the kinetics of two isomerization steps preceding HNH and RuvC cleavage. This model was derived in fitting Figure D, E and F simultaneously. The global fitting of all three experiments provides an accurate estimation for the rate constants for R loop formation and subsequent cleavage reactions. (D) Time dependence of HNH cleavage measured as in A. (E) Time dependence of RuvC cleavage, measured as in A. (F) R-loop formation was measured by mixing Cas9.gRNA (500 nM) with DNA with a 2AP label at position −9 on the non-target strand (100 nM) using stopped flow fluorescence method (Auto SF 120x, KinTek Corporation, Austin, TX.). The fluorescence increase as a function of time was biphasic defining two steps in R loop formation. The experiments including cleavage initiated by the simultaneous addition of DNA and Mg^2+^ for both HNH and RuvC cleavage were globally fit with the R loop formation measurement using the model shown in C.

To test this hypothesis, we directly measured the rate for R-loop formation using a stopped flow assay for DNA unwinding. We incorporated 2-aminopurine (2-AP) in the non-target strand at position −9 nt distal to the PAM (Figure S2). The opening of dsDNA upon R-loop formation leads to an increase in fluorescence because 2-AP fluorescence intensity is quenched by base stacking when it is within a dsDNA molecule but not in a disordered single-stranded state. The time dependent increase of the fluorescence signal was fit to a double exponential equation to obtain R-loop formation rates of 1 s^−1^ and 0.3 s^−1^ (Figure 3F). Notably, the trace shows two sequential increases in fluorescence, suggesting two sequential unwinding events. The stopped flow experiment was repeated using different concentrations of 2-AP labeled DNA, and the observed rates of the fast and slow phase were ~1 s^−1^ and 0.3 s^−1^ at all concentrations (Table S1). Interpretation of fluorescence signals is often ambiguous, especially when reactions are biphasic. We overcome these limitations by simultaneously fitting the data in Figures 3D-F, so that the changes in fluorescence are correlated with measureable chemical reactions for HNH and RuvC cleavage (Figure 3C). In developing a universal model, we also include the information from Figure 3B showing that if the DNA is allowed to equilibrate with the enzyme before adding Mg^2+^, cleavage is faster than R-loop formation.

The smooth lines shown in Figures 3D-F show the global fit derived according to the model shown in Figure 3C. The results show that when the reaction is initiated by adding DNA in the presence of Mg^2+^, the observed HNH cleavage is limited by the rate of the initial opening of double-stranded DNA as measured by the fast phase of the 2AP signal, which also accounts for the modest lag phase for RuvC cleavage (Figure 3E). The observed rate of RuvC cleavage is limited by the rate of the slower phase of the 2AP signal. These data demonstrate that the two cleavage events are not sequential, but can occur simultaneously. However, they are governed by two sequential DNA unwinding events that are reported by changes in 2AP fluorescence. Our global fitting of all three experiments (Figures 4D,E,F) supports a mechanism where the initial unwinding leads to HNH cleavage and a subsequent unwinding step leads to RuvC cleavage. We propose a model where R-loop formation is a two-step process. The HNH site is aligned for cleavage after the first step, while RuvC is aligned after the second step. In addition, we can account for the lower amplitude of the observed RuvC cleavage by the equilibrium constant for the second isomerization.

**Figure 4.**
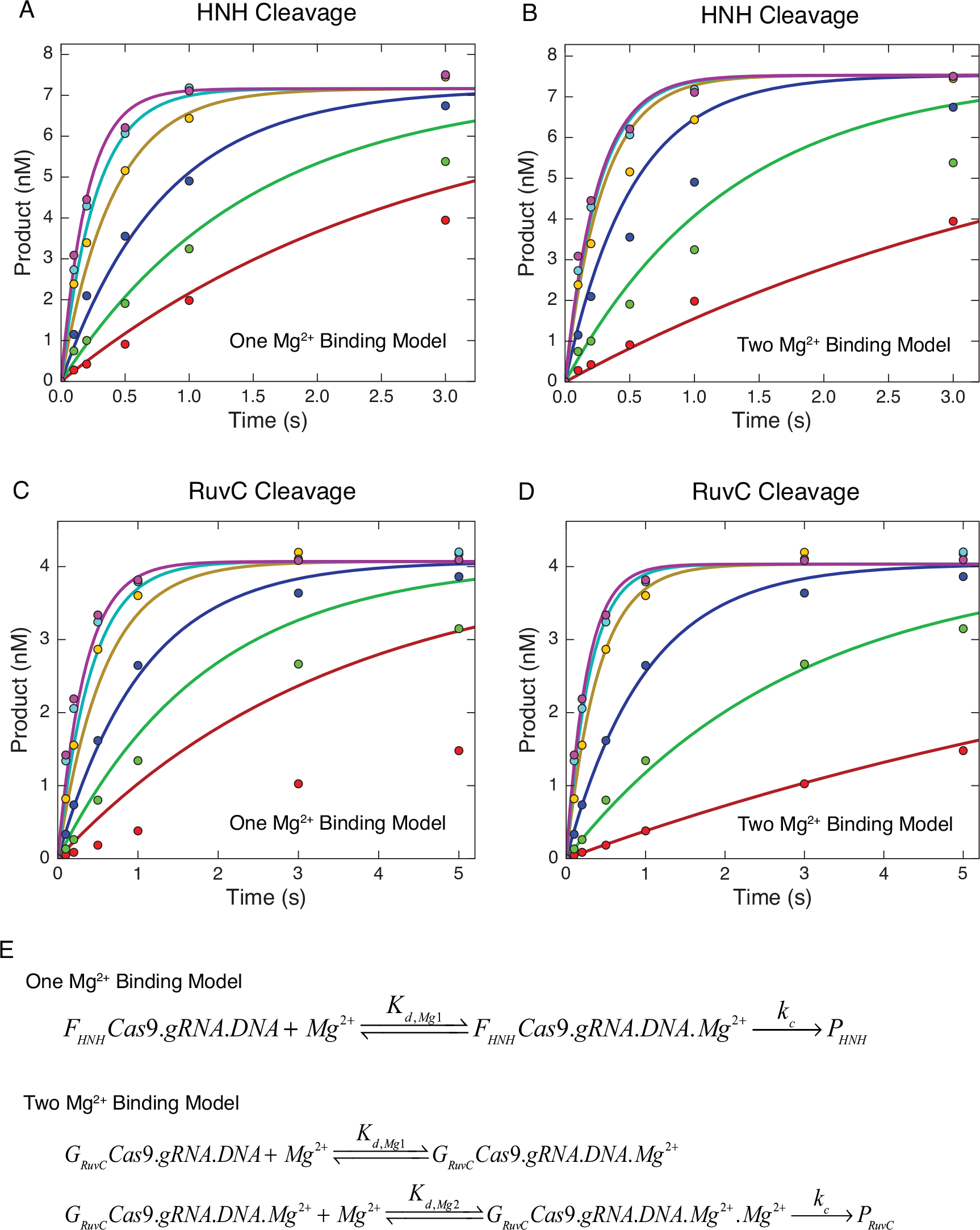
HNH and RuvC use one and two metal ion mechanisms, respectively. (A) Mg^2+^-dependent HNH cleavage experiment performed by premixing Cas9.gRNA (27.6 nM active site) with γ-^32^P labeled DNA (10 nM) in the absence of Mg^2+^ for 10 min. The reaction was initiated by addition of variable concentrations of free Mg^2+^ (0.5, 1, 2, 5, 10, 20 mM). The reaction was quenched by the addition of 0.5 M EDTA at different time points. The results were analyzed using a single Mg^2+^ ion binding model as shown in Figure 4E. (B) The same experiment performed in (A) analyzed using a two Mg^2+^ ion binding model as shown in Figure 4E. (C) Mg^2+^-dependent RuvC cleavage experiment performed as above analyzed using a one Mg^2+^ ion binding model. (D) The same experiment performed in (C) analyzed using a two Mg^2+^ ion binding model. (E) One Mg^2+^ binding model and two Mg^2+^ ion binding model.

### HNH and RuvC show different Mg^2+^ ion concentration dependence

We next examined the Mg^2+^ concentration dependence of Cas9 cleavage. Cas9.gRNA was premixed with DNA in the absence of Mg^2+^ to allow the formation of Cas9.gRNA.DNA complex. Then, the reaction was initiated by the addition of various concentrations of Mg^2+^. The amount of cleaved product as a function of time for both the HNH and RuvC domains was fit to a single exponential equation to define the cleavage rate at each concentration of Mg^2+^ (Figure 4A-D). The results show that both HNH cleavage and RuvC cleavage are Mg^2+^ dependent, as described previously (Jinek et al., 2012). The maximal cleavage rates (*k*_*c*_) for both HNH and RuvC nuclease domains were measured as >4 s^−1^ (Table 1), which suggests that the final cleavage step is fast in the presence of high concentrations of Mg^2+^. As described above, no lag phase was observed in either HNH or RuvC cleavage in the presence of various concentrations of Mg^2+^, indicating both HNH and RuvC cleavage occur immediately after Mg^2+^ binding. Thus, our results suggest that both HNH and RuvC cleavage occur simultaneously if DNA is bound, unwound, and all domains are positioned for cleavage before adding Mg^2+^ to simulate chemistry. Finally, our data for HNH and RuvC cleavage were fit to either a one Mg^2+^ binding model or two Mg^2+^ binding model (Figure 4E). These results show that HNH cleavage can be accurately fit with a one Mg^2+^ binding model (Figure 4A), while RuvC cleavage can be fit with a two Mg^2+^ binding model (Figure 4D). The apparent binding affinity (K_d,Mg1_) for one Mg^2+^ ion at the HNH active site is estimated as >5 mM (Table 1). The apparent binding affinities (K_d,Mg1_ and K_d,Mg2_) for two Mg^2+^ ions at RuvC active site are estimated as 1.6 mM and 6 mM, respectively (Table 1). These results are consistent with the one and two metal ions seen in the active sites of homologs of the HNH and RuvC domains, respectively.

**Table 1.** Kinetic Constants for Mg^2+^ Mediated Cas9 Cleavage

| Active sites | $k_c$<br>($\text{s}^{-1}$) | $K_{d,Mg1}$<br>(mM) | $K_{d,Mg2}$<br>(mM) |
| --- | --- | --- | --- |
| HNH | > 4 | > 5 | None |
| RuvC | 4.2<br>(3.9, 4.8) | 1.6<br>(0.4, 3) | 5.9<br>(3, 23) |

Note: 1. The $\chi^2$ threshold boundary was set at 0.9 to define the lower and upper limit for each parameter. The results are shown as (lower, upper limit).
2. $k_c$ represents the rate constants for cleavage step; $K_{d,Mg1}$ and $K_{d,Mg2}$ represent the apparent binding affinities for two $\text{Mg}^{2+}$ ions
3. The numbers in Mg1 and Mg2 do not indicate the order of binding.

### A complete kinetic framework for CRISPR-Cas9 activity

The experiments performed in our study define the constants in a complete kinetic framework of Cas9 (Figure 5). The result shows that Cas9.gRNA binds tightly to a perfectly matched DNA to form a Cas9.gRNA.DNA complex. It was previously reported that Cas9 undergoes a large conformational change upon RNA binding that activates the enzyme to form Cas9.gRNA, and DNA binding induces an additional rearrangement (Jinek et al., 2014). Here, we show that after this rearrangement most of Cas9.gRNA.DNA forms a reactive state, while a fraction of Cas9.gRNA.DNA goes into a nonproductive state. The transition from the nonproductive state to productive state is relatively slow. Following DNA binding, R-loop formation occurs. In this process, dsDNA is unwound with a rate constant of 1.6 s^−1^. For the HNH domain, cleavage can occur instantaneously if one Mg^2+^ is present. However, there is an additional step prior to cleavage required for RuvC domain activation. This step could either be further resolution of the R-loop or RuvC repositioning, or a combination of both. The rate for this additional step is 0.2 s^−1^. RuvC is then primed for cleavage if two Mg^2+^ are present in the active site. The maximal cleavage rate for both domains is >4 s^−1^, with R-loop formation being the rate-limiting step in catalysis.

**Figure 5.**
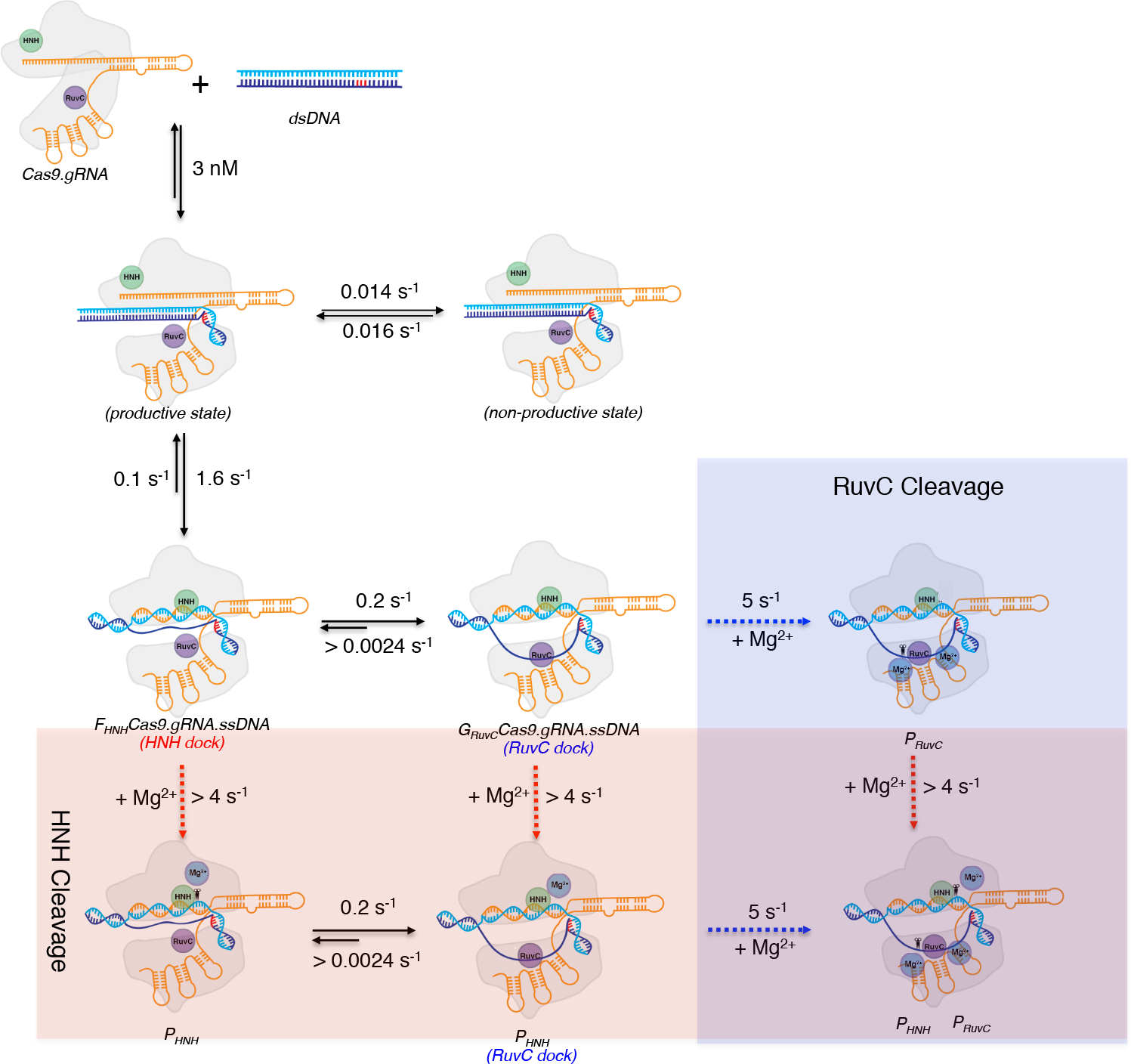
A complete kinetic framework for CRISPR-Cas9 activity. Our model for the kinetic pathway of Cas9 cleavage depicted using cartoons. The rate constants for each individual step including DNA binding, R-loop formation, Mg^2+^ ion binding, and HNH and RuvC cleavage were obtained from our experiments. The cartoon is color coded as follows: Cas9, outlined in light grey; HNH domain, green circle; RuvC domain, purple circle; Mg^2+^ ions, blue circles; sgRNA, orange; target strand, light blue; non-target strand, dark blue; and PAM, red. Scissors indicate cleavage. The transparent blue region indicates RuvC cleavage and the transparent red region indicates HNH cleavage.

### Discussion

Here, we show the first measurement for the active site concentration of Cas9.gRNA, the kinetics of R-loop formation and of HNH and RuvC cleavage. While recent single molecule measurements have provided some insights into the activity of Cas9, we use measurements of the actual chemistry of the enzyme to determine the kinetic pathway. Moreover, by simultaneously fitting data from fluorescence and rapid quench experiments, we show how the measured fluorescence changes relate to the chemical reaction. It should be noted that the rate of the chemical reaction is relatively fast, so single molecule measurements reflect signals arising after the chemical reaction.

The measurement of active site concentration is critical because it provides the basis for accurately measuring kinetic parameters in the cleavage pathway. The active site concentration of Cas9.gRNA complex in our study was 28 nM using nominal concentration of 100 nM Cas9.gRNA (1:1 ratio) based on absorbance measurements. Most in vitro studies of Cas9 activity use a 10-fold excess of Cas9.gRNA over substrate, possibly reflecting the low active site concentrations in many protein preparations. Future studies will be needed to establish if the low active site concentration is because of poor protein-stability, inefficient protein-folding during or after purification, or systematic errors in absorbance measurements needed to estimate protein concentration. However, we do recommend that our simple protocol (Figure 2A) become standard practice to establish active site concentration.

EM and x-ray crystallographic studies of Cas9 have previously shown Cas9 undergoes major structural rearrangements upon sgRNA binding (Jinek et al., 2014, Nishimasu et al., 2014, Jiang et al., 2015), creating extensive contacts between the RNA and enzyme. In the presence of 100 nM Cas9.gRNA (1:1 ratio), ^~^85% of the target strand is cleaved by HNH after 30s, while only ~35% of the non-targeted strand was cleaved by RuvC (Figure S1). Increasing concentrations of gRNA does not increase the amplitude of the product formation by the HNH domain buts lead to a significant increase in amplitude of product formation by the RuvC domain (Figure S1). The increase in RuvC cleavage in a guide RNA-dependent manner is still a mystery. We speculate that only a fraction of the *in vitro* transcribed and re-folded gRNA is productively interacting with the Cas9 enzyme. Incompletely folded gRNA would still likely have the single stranded seed available for target recognition and HNH-mediated cleavage, but improperly folded stem loops would not create the contacts required for the activation of the RuvC domain.

A major finding of this study is that R-loop formation is rate-limiting and is followed by rapid HNH and RuvC cleavage events. Moreover, the kinetics demonstrate that HNH and RuvC cleavage events are not sequential, but rather can occur simultaneously. Prior to this work, the rate-limiting step in Cas9-mediated cleavage had not been identified. Here, we show that R-loop formation is the rate-limiting step. Interestingly, this finding suggests that most studies measuring Cas9 cleavage are actually measuring the rate of R-loop formation. A previous study gave and observed rate for Cas9 cleavage of 9 min^−1^ (0.15s^−1^) (Sternberg et al., 2014), which is equivalent to our measured second step in the R-loop formation rate. The actual chemistry of cleavage is much faster. Both HNH and RuvC cleavage occur simultaneously and at a rate ~ 4 s^−1^ under conditions where DNA has been pre-incubated prior to addition of Mg^2+^. When the reaction is initiated by DNA binding in the presence of Mg^2+^, HNH and RuvC cleavages are limited by sequential isomerization reactions, which may make it appear that the two cleavage reactions are sequential under some assay conditions. That is, we did identify a brief lag phase when RuvC-mediated cleavage was initiated by the simultaneous addition of DNA and Mg^2+^, suggesting differences between how the HNH and RuvC domains engage with the R-loop. The initial fast phase (1s^−1^) of R-loop formation corresponds to HNH recognition and cleavage. This step is followed by a slower phase (0.2s^−1^), which is correlated with the lag phase in RuvC recognition and cleavage. This appears to be due to a second step in R-loop formation (i.e. further unwinding) as reported here, which could represent the postulated conformational checkpoint between the HNH and RuvC domains described previously (Sternberg et al., 2015, Dagdas et al., 2017). We propose that in addition to HNH recognition, there is another step involved in DNA unwinding that is required for processing, irrespective of mismatches. This model would be consistent with the structure of Cas9 bound to an R-loop, where the non-target strand is disordered, undergoes dramatic kinking in several regions, and enters the RuvC domain through a tight tunnel (Jiang et al., 2016). Therefore, it is conceivable that additional changes in the non-target strand conformation are required before cleavage. A different single molecule FRET study identified sub-conformations of the RNA–DNA heteroduplex during R-loop expansion (Lim et al., 2016), which may be explained by our model. While this report also suggested that the rate of R-loop formation was 2 orders of magnitude faster than the cleavage rate, we unambiguously identify R-loop formation as the rate-limiting step in Cas9-mediated cleavage. Our model implies an induced-fit model for Cas9 selectivity in which the initial binding of DNA is followed by a dramatic, rate-limiting and largely irreversible isomerization of the enzyme-DNA complex leading to faster catalysis. This follows the paradigm for DNA polymerase fidelity (Kellinger and Johnson, 2010), suggesting that the DNA dissociation rate (reversal of R-loop formation) may be the major determinant of the specificity constant (how efficiently an enzyme converts substrates into products).

Cas9 is a single turnover enzyme that binds perfectly matched targets with a correct PAM tightly. It dissociates very slowly from DNA with a rate of 0.0024 s^−1^ prior to cleavage. At long time scales (>30 min), the transition from the nonproductive state to the productive state drives cleavage to completion and makes the DNA binding appear to be much tighter because the reaction is dominated by the nearly irreversible product release. When used in *in vivo* cellular applications, other host factors must be required for removal of the Cas9 enzyme and repair of the DSB. It is tempting to hypothesize that the DSB repair factors themselves are responsible for removing the Cas9 protein from DNA.

The nonproductive state of Cas9 is intriguing. This state differs from inactive enzyme because the slow transition from the nonproductive state to the productive state allows cleavage to come to completion. The effects of the DNA trap on the nonproductive state can be observed after 10s (Figure 2A), suggesting that the off-rate of DNA in this state is much faster than the off-rate measured for the productive state (0.0024s^−1^). In addition, the rate of conversion from nonproductive to productive state was calculated as 0.016s^−1^ based on our global fitting of all three experiments, which indicates that DNA can dissociate much faster from the nonproductive than the productive state. Previous studies show that partial R-loop formation results in the reduction of cleavage by Cas9.gRNA (Lim et al., 2016), and our data show the nonproductive state is also inefficient. Taken together, we hypothesize that this nonproductive state contains an incomplete or partially opened R-loop that can collapse and be ejected from Cas9.gRNA. A recent study using FRET showed that the HNH domain could possibly exist in an intermediate state (Dagdas et al., 2017). Future experiments are required to determine if the state they observed is the same as the one identified here.

While many comparisons have been performed between the HNH and RuvC domains with homologous structures, the precise metal-ion requirement for the cleavage mechanism has remained elusive. Because Mg^2+^ ions are normally excluded for crystallographic studies of Cas9 bound to substrates to prevent cleavage, it is challenging to directly capture Mg^2+^ ions in the HNH and RuvC domains. Our results also show that the apparent binding affinity of Mg^2+^ ions is weak, suggesting that most experiments using 2–5 mM of free Mg^2+^ do not actually saturate the active sites. Additionally, our results show that the RuvC domain uses a two-metal ion mechanism for catalysis. Our measurements also strongly support the one-metal ion mechanism suggested for the HNH domain.

Here, we have defined the complete kinetic pathway for Cas9 cleavage. Global fitting of experiments together allow us to obtain the kinetic parameters for each step in the pathway. Our results show that binding of DNA is very tight (K_d_ < 3 nM), most likely mediated by specific interactions between Cas9 and the PAM and hybridization of the sgRNA with the target strand of the DNA target. Cas9.gRNA complex exists in both a productive and minor nonproductive state upon DNA binding. The productive state leads to the R-loop formation and the opening of dsDNA with a rate of 1s^−1^. HNH cleavage occurs immediately after the opening of the two DNA strands, while RuvC requires an additional activation step prior to cleavage. Future experiments will be required to elucidate the identity of the nonproductive state. It will also be interesting to test the effect of mismatches within target DNAs and high-fidelity versions of Cas9 using our new kinetic scheme.

### Contact for Reagent and Resource Sharing

Further information and requests for resources and reagents should be directed to and will be fulfilled by the Lead Contact, David W. Taylor.

### Experimental Procedures

#### Expression and purification of Cas9

Plasmid pMJ806 containing *Streptococcus pyogenes* Cas9 was obtained from Addgene (Cambridge, MA). For bacterial expression of the recombinant Cas9, pMJ806 plasmid was transformed into BL21-Rosetta 2 (DE3)-competent cells (Millipore). The *E. coli* cells were cultured at 37°C in LB medium (containing 50 mg/l kanamycin) until the OD600 reached to 0.5-0.8, and then Cas9 expression was induced by the addition of 0.25 mM isopropyl-β-D-thiogalactopyranoside (IPTG) for 20 hours at 18°C. The His_6_-MBP tagged Cas9 was purified by a combination of affinity and size exclusion chromatography, essentially as described previously (Jinek et al., 2012) with the following modifications. Briefly, bacterial cells were lysed by sonication in buffer containing 20 mM Tris-HCl pH 8.0, 500mM NaCl, 1mM phenylmethanesulfonyl fluoride (PMSF) and Piece protease inhibitor cocktail (Thermo Scientific). Clarified lysate was applied to HisPur^TM^ Ni-NTA resin (Thermo Scientific) and the resin was washed extensively with buffer containing 20 mM Tris-HCl pH 8.0, 500mM NaCl, 1mM PMSF, 10mM Imidazole and 10% glycerol. The bound protein was eluted in 20 mM Tris-HCl pH 8.0, 250mM NaCl, 1mM PMSF, 10% glycerol and 100mM Imidazole. The eluted protein was dialyzed and His_6_-MBP tag removed by addition of TEV protease to the dialysis tubing (Spectrumlabs) against buffer containing 20mM HEPES pH 7.5, 150mM KCl, 10% glycerol. Cas9 was further purified by size exclusion chromatography on a Superdex 200g 16/600 in 20mM HEPES pH 7.5, 150mM KCl. Cas9 protein was concentrated to ~5mg/ml and stored at −80°C. Protein concentration was calculated using extinction coefficients of 120,450M^−1^cm^−1^ at 280 nm for Cas9.

### *In vitro* sgRNA transcription and refolding

The primers used as templates for sgRNA transcription and sequence of the sgRNA are listed in Key Resources Table. Both complementary oligonucleotides at equal molar concentration were mixed in 10mM Tris-HCl pH 8.0, 50mM NaCl, 1mM EDTA and heated to 95°C for 5 minutes, then slowly cooled to room temperature. The sgRNA was *in vitro* transcribed using the HiScribe Qiuk T7 RNA synthesis kit (New England Biolab) following the manufacturer’s protocol. The transcribed sgRNA was purified using a PureLink column (Thermo Scientific). The purified sgRNA was refolded by heating to 95°C and then slowly cooling down to room temperature in 10mM Tris-HCl pH 8.0, 50mM NaCl, 1mM EDTA.

### DNA duplex formation and probe labeling

55-nt DNA duplexes were prepared from unmodified and 2-aminopurin (2-AP)-labeled DNA oligonucleotides synthesized and PAGE gel purified by Integrated DNA Technologies. The synthesized oligonucleotides and the position of 2-AP labeling are listed in Star Resources Table. The DNA duplex used for RuvC cleavage assays was prepared by γ ^32^P labeling the nontarget strand before annealing with the cold complementary strand at a 1:1.15 molar ratio. The DNA duplex used for HNH cleavage assays was prepared by γ ^32^P labeling the target strand before annealing with cold nontarget strand at a 1:1.15 molar ratio.

### Quench flow kinetic assay

Quench flow experiments were performed by rapidly mixing cas9-gRNA complex (1:1 ratio) with γ ^32^P labeled DNA substrate at 37° using KinTek RQF-3 instrument (KinTek Corporation, Austin, TX). The reaction was stopped by the addition of 0.5 M EDTA at varying time points. Products were collected and separated on a 15% denaturing PAGE (acrylamide (1:19 bisacrylamide), 7M Urea) gel and quantified using a Typhoon scanner with ImageQuant 6.0 software (Molecular^®^ Dynamics).

### Stopped flow kinetic assay

Stopped flow experiments were performed by rapidly mixing cas9-gRNA complex (1:1 ratio) with 2AP-labeled DNA substrate at 37° using an AutoSF-120 stopped-flow instrument (KinTek Corporation, Austin, TX). Samples were excited at 315 nm, and the time-dependent fluorescence change was monitored at 340 nm using a single band-pass filter with a 26-nm bandwidth (Semrock).

### Global analysis of kinetic data

The kinetic data defining Cas9 cleavage were globally fit to the models shown in schemes 1-5 by *KinTek Explorer* software (KinTek Corporation. Austin, TX) to obtain rate constants shown in each model. FitSpace confidence contour analysis was performed to define the lower and upper limits for each kinetic parameter.

### Active site titration assay

The active site titration experiment was performed by mixing fixed concentration (100 nM) of Cas9.gRNA (1:1 ratio) with variable concentrations of γ ^32^P labeled DNA at time scales of 30s and 30min. Data were then analyzed by non-linear regression using the program GraFit 5 (Erithacus Software, Surrey, UK). The data was then fit to quadratic equation to obtain the apparent binding affinity (*K*_*d,app*_) of DNA and the active site concentrations (*E*_*0*_) of Cas9.gRNA.

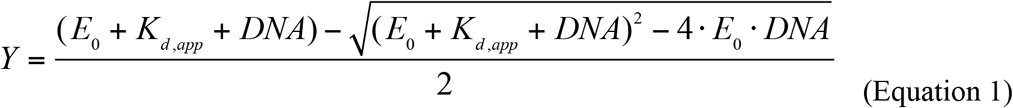

Where *Y* represents cleavage product at 30s or 30min, *E*_*0*_ represents the active site concentration of Cas9, DNA represents the concentrations of DNA that were used in the assay and *K*_*d,app*_ represents the apparent binding affinity of DNA.

### Calculation of DNA binding affinity

In the simple binding model of DNA:

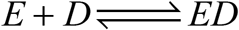

The binding affinity of DNA could be calculated using the equation:

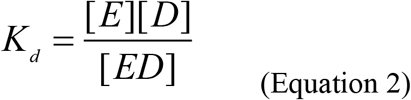

Where K_d_ represents the binding affinity of DNA; E represents the enzyme concentration calculated by subtracting ED concentration from initial enzyme concentration; D represents DNA concentration calculated by subtracting ED concentration from initial DNA concentration; and ED represents the final product concentration.

## Author Contributions

S.G. performed kinetic studies. H.H.Y. purified and reconstituted Cas9.gRNA complexes and assisted with kinetic studies. S.G. and K.A.J. analyzed the kinetic data. S.G., K.A.J. and D.W.T. interpreted the results and wrote the manuscript. K.A.J. and D.W.T. conceived experiments, supervised research, and secured funding for the project.

## Acknowledgements

We thank E. Schwartz and N. Sun for assistance with protein purification; I. Finkelstein and R. Russell for critical reading of the manuscript; and members of the Johnson and Taylor laboratories for helpful discussion. This work was supported in part by Welch Foundation Grants F-1604 (to K.A.J.) and F-1938 (to D.W.T.). D.W.T is a CPRIT Scholar supported by the Cancer Prevention and Research Institute of Texas (RR160088). K.A.J. is President of KinTek Corp., which provided the stopped-flow and chemical-quench-flow instruments and the KinTek Explorer software used in this study.

